# Deep neural generation of neuronal spikes

**DOI:** 10.1101/2023.03.05.531237

**Authors:** Ryota Nakajima, Arata Shirakami, Hayato Tsumura, Kouki Matsuda, Eita Nakamura, Masanori Shimono

## Abstract

In the brain, many regions work in a network-like association, yet it is not known how durable these associations are in terms of activity and could survive without structural connections. To assess the association or similarity between brain regions with a new “generating” approach, this study evaluated the similarity of activities of neurons at the cellular level within each region after disconnecting between regions. To this end, a multi-layer LSTM (Long-Short Term Memory) model was used. Surprisingly, the results revealed that generation of activity from one region to other regions that had been disconnected was possible with similar reproduction accuracy as generation between the same regions in many cases. Notably, not only firing rates but also synchronization of firing between neuron pairs, which is often used as neuronal representations, could be reproduced with considerable precision. Additionally, their accuracies were associated with the relative distance between brain regions and the strength of the structural connections that initially connected them. This outcome not only enables us to look into principles in neuroscience based on the potential to generate new informative data, but also creates neural activity that has not been measured in adequate amounts and could potentially lead to reduced animal experiments.

## 1. Introduction

### 1-1. Researches on multicellular neural activity

In the nervous system, a large number of neurons are repeatedly firing as they interact with each other. This scene has been likened to a symphony of complex spikes [Varela, 1995; Buzsaki, Draguhn, 2004; Engel et al., 2002]. When measuring neural activity, it is customary for electrophysiologists to discern from the sound of spikes whether the measurement points are in contact with active neurons, and then to be satisfied or disheartened.

It is also known that certain long-time correlations exist in the spike time series of neural activity [He, 2012; Martinello et al., 2017; Safonov et al., 2010; Fagerholm et al., 2016], indicating that an activity state of a neuron population at a time already has some information about its future state. The state of whether or not an individual neuron fires is essentially maintained as an interdependent relationship among multiple neurons, rather than maintained individually for inherent activity mode of each neuron.

The number of neurons that can be measured simultaneously is growing every year [Brown et al., 2004; Stevenson, Kording, 2011]. Thanks to these advances, we now have the capability afforded by recording technology necessary to reliably verify how accurately we can generate the future activity of individual neurons from past activity of the neuron population.

### 1-2. Time series data-generation

When we consider a symphony of neural activity (i.e., a coordinated flow in time) as music, we notice an interesting technical diversion. Polyphonic music includes multiple musical notes sounding simultaneously as found in piano music and ensemble music which can be regarded as time series data similar to multicellular spikes data. We are mapping the pitches of musical notes to the neurons and the onset time to the time of firing. The existence of co-occurrence relationships and long-time correlations between specific pitches is similar for music data.

Attempts to automatically generate music have been made since the 1950s [Ames 1987]. Many have also utilized Artificial Neural Networks (ANNs) for music generation [Todd 1989; Eck, Schmidhuber, 2002]. Along with recent advances in data analysis techniques using ANNs, music generation techniques based on ANNs have been significantly improved.

For example, a method was developed to solve the gradient loss problem of recurrent neural networks (RNNs) [Hochreiter, Schmidhuber, 1997], and a long short-term memory (LSTM) network, a type of RNN that considers the effects of short- and long-term memories, has been applied [Boulanger-Lewandowski et al., 2012]. A multi-layer LSTM network is an ANN with multiple layers of LSTM units between the input and output layers. Recently developed techniques of ANNs, such as the Generative Adversarial Network (GAN) and Transformer, have also been applied to music generation, making it possible to generate music data that better approximate real data with long-time correlations and dependencies among musical notes [Huang et al., 2018; Dong et al., 2017].

We can expect that it is also possible to generate spike data with properties similar to those of real-world music.

Despite the fact that ANNs are based on nervous system motifs, surprisingly, they have rarely been applied to neural activity spike generation [Gers et al. 2002] and have not yet been fully deployed. Technology to automatically generate spikes by applying Artificial Neural Networks for analyzing neural spike data has long remained unexplored.

### 1-3. A brief research history of neural spontaneous activity

The nervous system is active even in the absence of external stimuli. Such neural activity is called spontaneous activity. For a long time, neural activity has been measured while animals undertake any tasks and the neural activities have been evaluated in correlation with the task. In fact, more than 80% of the energy in the brain is expended in the task-free state, and spontaneous activity consumes most of the energy of neural activity [Raichle, 2015].

As we will discuss later, it is also known that stimulus-induced activity is fundamentally rooted in the state of preceding spontaneous activity.

In the past, when many neurons could not be measured simultaneously, temporal changes in the activity of individual neurons were regarded only as classical stochastic activity. However, recent measurements have shown that spontaneous activity is also considered to retain a causal relationship between activities with a degree of inevitability [Kaminski et al., 2001; Quinn et al., 2011].

On a macroscopic (anatomical) scale, spontaneous activity has been observed to produce specific patterns throughout the brain. A typical example is the default mode network, a pattern of activity that is inversely correlated with presentations of external stimuli [Raichle et al.]. It is also clear that there are multiple other modes in the macroscopic spontaneous activity patterns [Power et al., 2014].

This massive amount of research on spontaneous activity on a macroscopic scale forms a huge research field that continues to this day. The spontaneous functional activity patterns can also be systematically interpreted by comparing them to structural wiring [Jonston et al. 2008; Honey et al. 2010; Goni et al. 2014; Rosenthal et al. 2018].

The measurement and analysis of spontaneous activity of neurons at the microscale have been pursued both in vitro and in vivo. Classically, the firing timings of individual neurons have been quantified as a deviation from the Poisson point process generated when we regard them as a random time series [Heeger, 2000; Kass, Ventura, 2001]. Randomness and simple repetitive patterns have also been assumed in the activity patterns of multiple neurons.

Recent studies have begun to capture the presence of complex, but non-random rules within these patterns. One of pioneering studies, using real-time optical imaging, revealed the patterns of spontaneous activity observed when multiple neuronal activities are measured simultaneously. For example, in the rodent visual cortex, spontaneous activity was found to intrinsically exhibit variations of spatial patterns in evoked activity even before stimulus presentation [Arieli et al., 1996]. The same research team also demonstrated that activity patterns obtained from optical imaging time-locked to the firing timing of single neurons show significant similarity to patterns time-locked to evoked activity [Tsuodyks et al., 1999].

Such interactions between multiple neurons have sequential patterns caused by a series of inevitable interactions that are thought to be mediated by synaptic connections between neurons.

The connections of quantified causal interaction between neurons drawn as arrows is called effective connectivity. Much work has also been done to reconstruct structural wiring as effective connectivity reconstructed from neural activity [Stetter et al. 2012; De Blasi et al. 2019; Gu, et al. 2019]. It has also been pointed out that networks reconstructed from neural activity are closely related to stimulus-dependent evoked activity of neurons [Bock et al., 2011].

### 1-4. Topics conducted in this research

The main goal of this study is to generate neuronal spike data using one of the techniques described in section 1-2 that can capture causal interactions between neurons. Beyond the naive methodology of using correlations between spike’s data, we evaluated the similarities and differences between real and generated neural activities in terms of predicting future neural spikes.

The overview of the entire data processing flow in this study is summarized in Figure 1.

**Figure 1.**
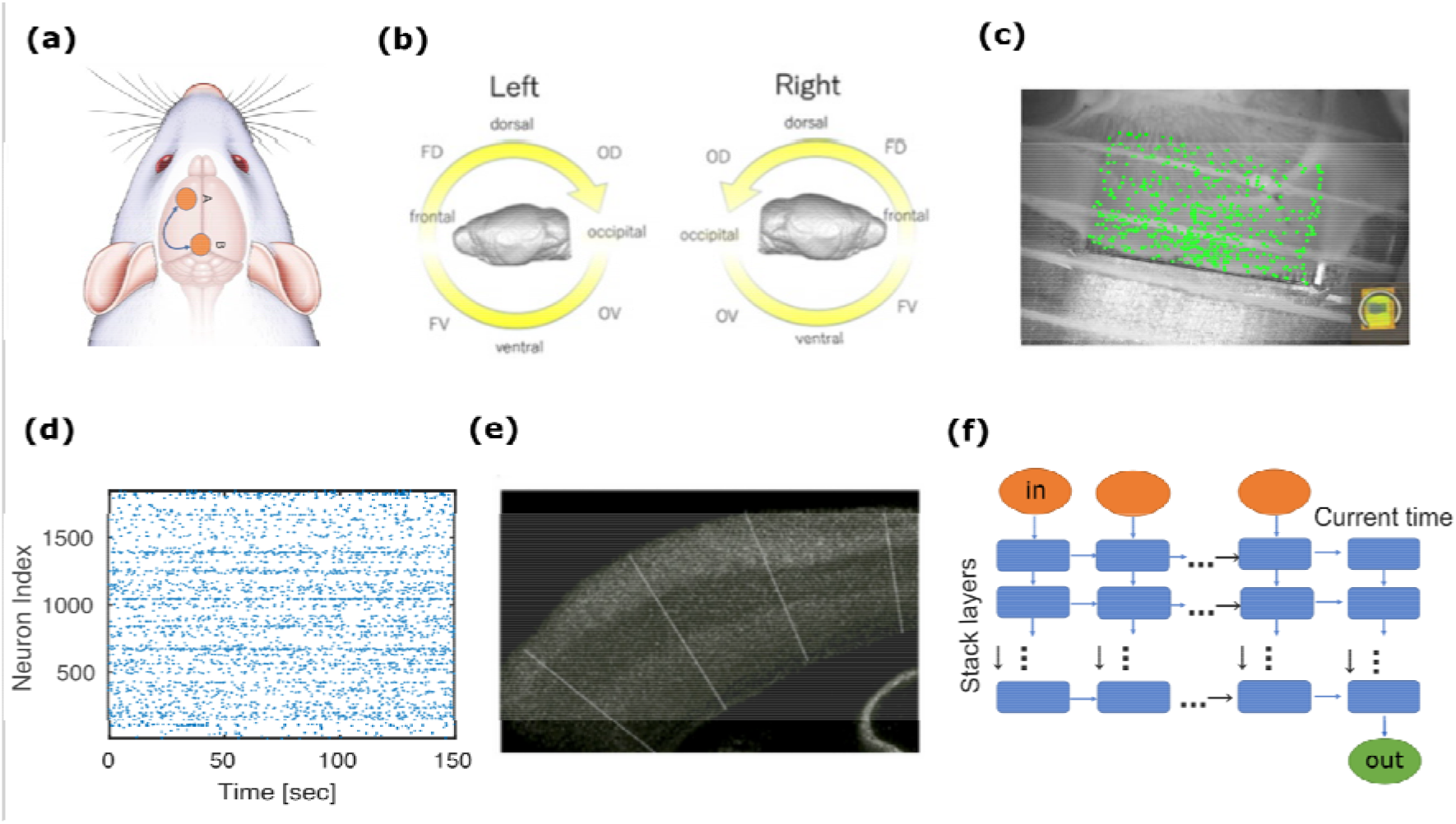
The workflow of this study: (a) We prepared brain slices from individual regions, as illustrated here with orange circles. We used two of these regions for the training step and the generation step, respectively. (b) Prior to this, we classified the cortical regions into 16 groups (see Table 1 for their short names). Pairs of regions, like the examples in panel (a), were selected from those 16 groups. (c) We measured neuronal activity from hundreds of neurons in each region with a multi-electrode device. (d) An example of a spike train obtained from one of the measurements. The horizontal axis is time [sec] and the vertical axis is the index of neurons. The timing at which a certain neuron fires is indicated by a dot. This diagram is called a raster plot in neuroscience. (e) We used stained images to extract only neuron groups in cortical areas and then divided the neuron groups with lines orthogonal to the cortex so that only 128 cells were included in each data set. (f) Such spike sequence, binary data, is input to the Multi-layer LSTM model to predict one step ahead after learning from the past data. The horizontal axis is time [ms], and the input vector is a binary vector.

This study targeted the analysis of electrical activity of multiple neurons in the mouse cortex, mainly neocortex, measured with a Multi-Electrode Array (MEA) (fig1-c,d). The neocortex consists of one to six layers, numbered from the surface in the direction of the depths. In each brain region, neurons were selected by sliding a section orthogonal to the cortical surface along the cortical surface so that all 1-6 layers were included, and a total of 128 cells were selected and collected into each regional data (fig1-e, Refer method 4-1 in detail).

The neural activity recorded with an MEA is called spikes as mentioned before, which is represented as binary data, where elements with 1 describe firing timings.

In individual analysis, we prepared a pair of training and test spike data from two datasets. Training data is used to optimize the internal parameters of the multi-layer LSTM model, enabling the best prediction within the training data. This is called the training process. After training is finished (fig1-f), we use the trained network to generate new spike data and compare it with the test data (fig1-a) (Refer to method 4-3-2, 4-3-2 for more details). This is called the generating process. In the generating process, we generated one step future neuronal spikes with hypothesizing that we know all neuronal spike sequences.

The test data is also sometimes referred to as target data because this data is the target of the generation process. The training and test data were respectively obtained from one of the 16 regions of the neocortex (fig1-b, Table 1) [Matsuda et al., 2022].

**Table1:**
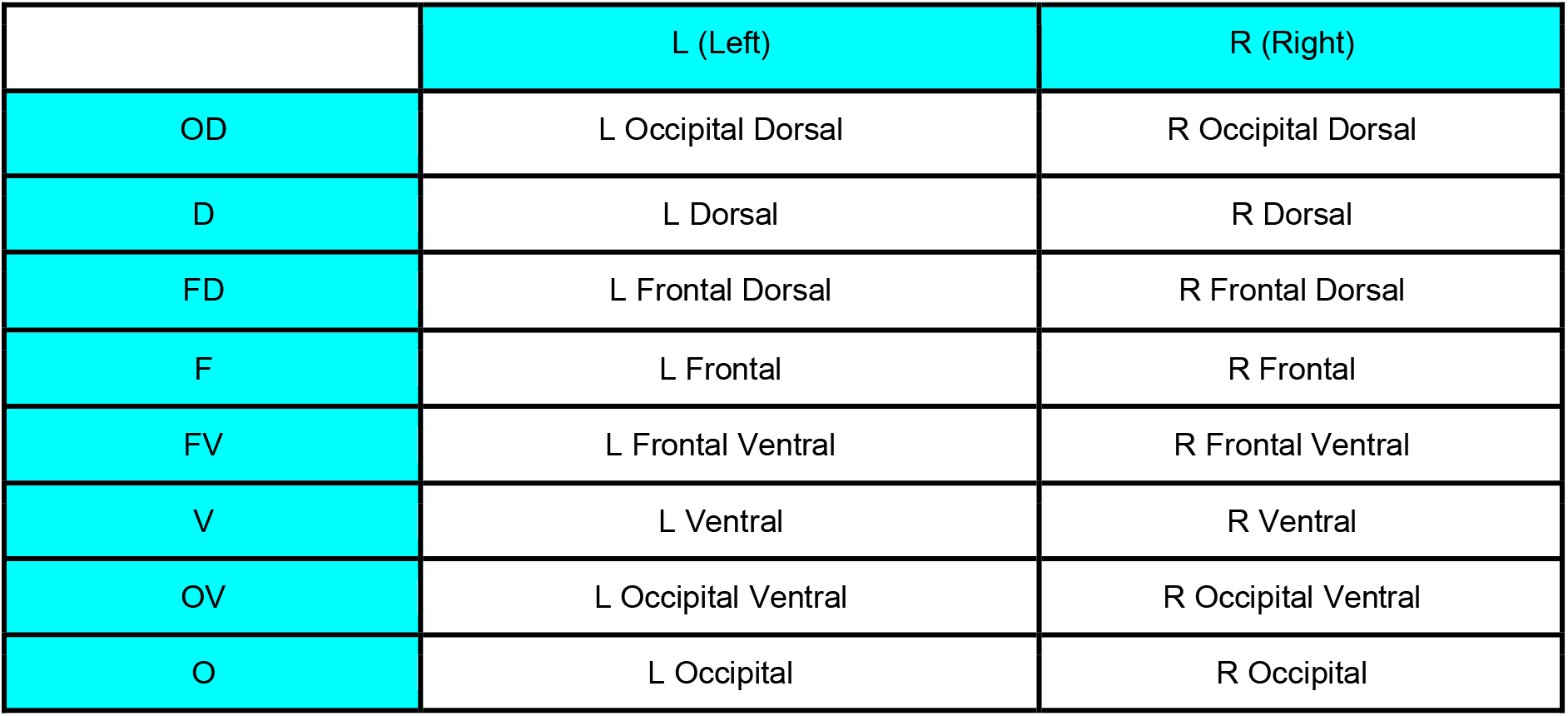
Definition of 16 regional groups: Right and left hemispheres include 8 groups, respectively, and expressed as L or R at the beginnings of individual names. The name is followed by combinations of O, D, F and V expressing abbreviations of Occipital, Dorsal, Frontal and Ventral. (Refer to the supplemental material for detailed locations of the slices used for the 16 area groups.)

The generated data were evaluated using the firing rate and the Synchronization score (Method 4-3-3). It is important to note that the ability to generate a highly predictive time series means that new future neural activity can be generated by extending time from existing spike data. We think that the above is important because it means that new time series data can be obtained without the need for new experiments, leading to fewer physiological experiments in the future.

In particular, when a time series generated by a model trained using data in one brain region, original data, is evaluated with test data obtained in another brain region, we can systematically investigate how the generation performance depends on the relative “closeness” of the two brain regions. Not only were we concerned about the geometric distance, but we also were concerned about connection strength of structural wiring through white matter fibers between the source and target brain regions.

## 2. Results

### 2-1. Training process

The internal parameters of the Multi-layer LSTM model were optimized to minimize the prediction error for approximately 17 minutes of training data.

As described in Method 4-3-1, based on the result of a preliminary study testing various parameterizations, we use a Multi-layer LSTM network with three hidden layers, each with 128 LSTM units. We used a focal loss function to quantify the prediction error [Refer method 4-3-3]. The number of epochs for training was set to 350. This was chosen based on the fact that, although even after the value of the loss function, had converged at about 25-100 epochs, the precision of the firing rate and the reproducibility of synchronous firing improved [Fig.2]. The Adam algorithm was used for optimization [Kingma, Ba, 2014].

**Figure 2.**
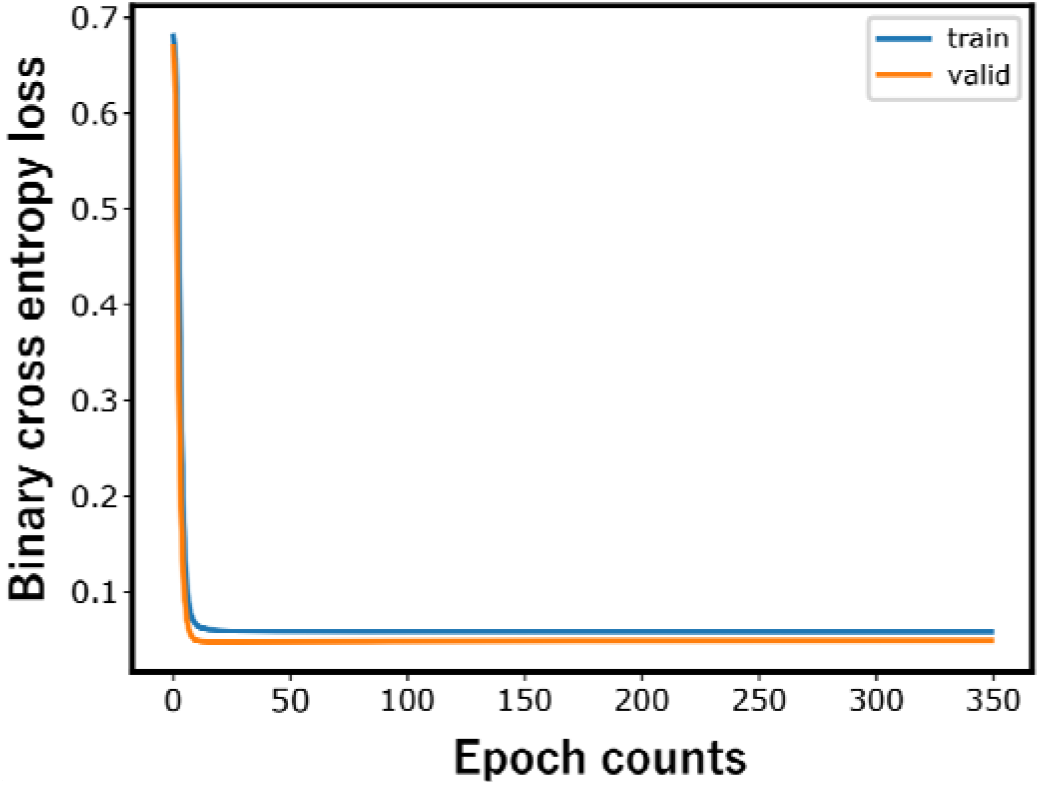
Learning procedure: The figure depicts how the loss on the training data and the loss on the validation data decreases as the multi-layer LSTM model is trained. A decrease in loss indicates that the training of the Multi-layer LSTM has progressed. The loss, common to both data, decreases sharply at 25-100 epochs.

### 2-2. Generation with a same region for source and target regions

The neocortex was divided into 16 regional groups, with two datasets per regional group. Formally, the first 16 data sets are collectively named dataset 1 and the remaining data sets are named dataset 2. In the following sections, we observe the results of evaluating the data generated as a result of the training in various cases. Then, we present the average of the results obtained for these two data sets. The results presented in what follows are confirmed to be reproducible between the two data sets.

In this section, we first performed generation through the multi-layer LSTM model by dividing the data given by the same group of brain regions in each dataset into the first half and the second half on the time axis, and preparing them as training data and test data, respectively [Fig. 3-(a,b)]

**Figure 3.**
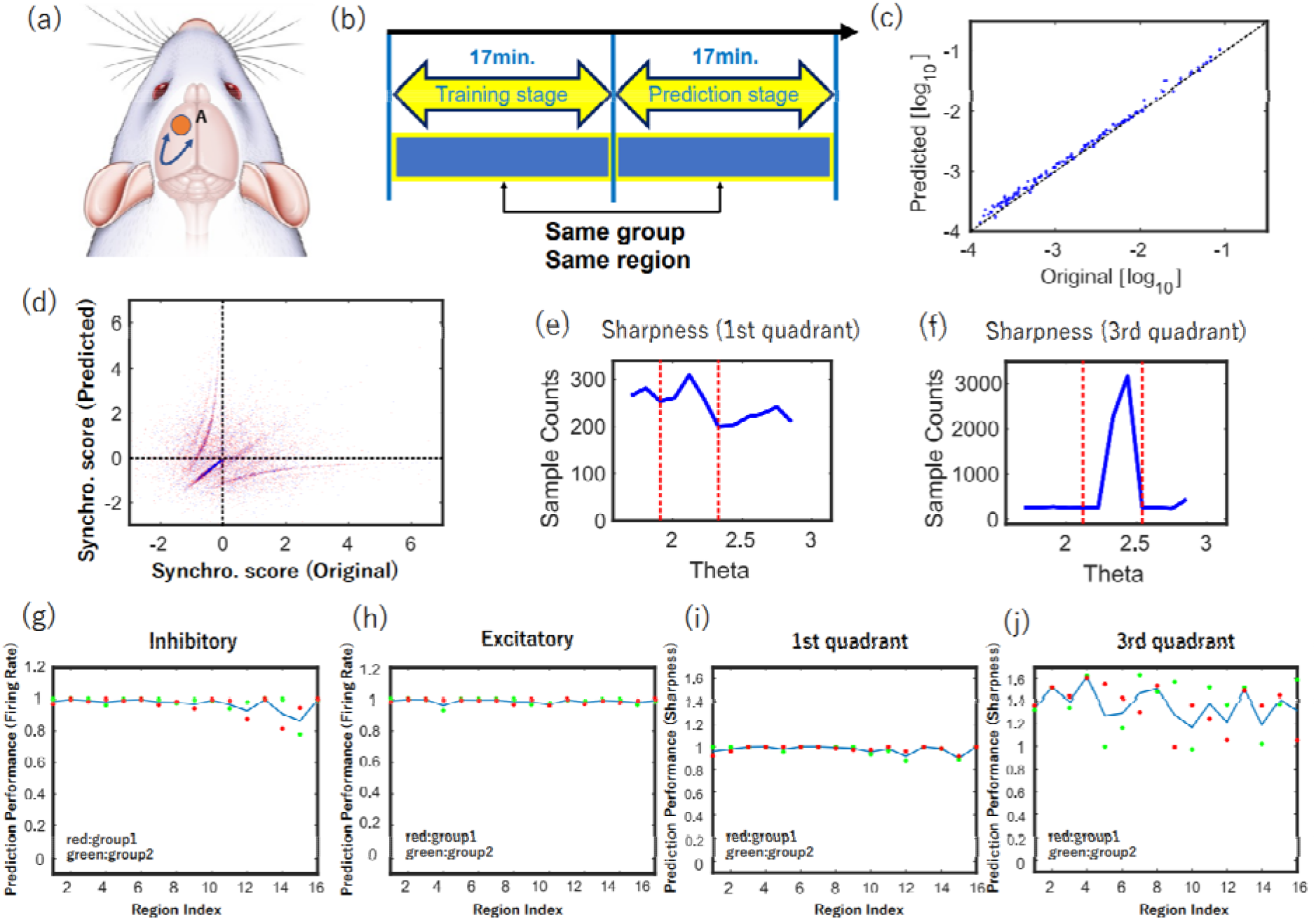
(a) Both training data and prediction, generation, data are prepared from the same region in this evaluation. (b) In the case of example (a), both training data (first 17 minutes) and test data (second 17 minutes) of the same time series are obtained from area A. (c) This panel shows the result of predicting the firing rate in this case, where the x-axis is the firing rate in the original data and the y-axis is the firing rate in the data generated by training Multilayer LSTM. (d) This panel shows the result of predicting synchronization score. Again, the x-axis is the synchronization score in the original data, and the y-axis is the synchronization score in the generated data. This data was expressed in r-θ rotational coordinates, and a histogram of the number of data in 0-π/2 with respect to θ, or in the first quadrant, was drawn in panel(e). Finally, panel(f) is the histogram of the number of data in π-3/2π with respect to θ, or the third quadrant. In particular, if the output is coming from inhibitory cells, it is distributed in the third quadrant. The sharpness of the peaks in these histograms (e) and (f) was evaluated by sharpness [Refer 4-3-4]. (g) Correlations between expected and true values of firing rates in the inhibitory neurons are plotted for every 16 regions. The two points for every group of regions correspond to the two data sets, and the line is the averaged value between the two data sets. The meaning of the points and lines is the same for (h)-(j). (h) Correlations between expected and true values of firing rates in excitatory neurons are plotted in the same way as in (g). (i) shows correlations between expected and true values of synchronization score in the first quadrant for every 16 regions. (j) shows correlations between expected and truth values of synchronization score in the third quadrant.

Multi-layer LSTM inputs time series data of 128 cells in the past and outputs information on whether 128 neurons are active in the future (fig. 1-f). Multi-layer LSTM was trained by swiping data in the first half of the time from time 0 to 17 min (Refer in more detail to the method sections 4-3-2 about Multi-layer LSTM). In the second half, the learning process is stopped, and the data is swiped from 17 min to 34 min to evaluate how well the rules learned in the first half can be used to predict future activity states. In other words, the similarity between the first half of the data and the second half of the data is evaluated through the data generation performance.

The accuracy of how well the generated time series reproduced the statistical properties of the actual data was evaluated as the average of dataset 1 and dataset 2. The accuracy was evaluated using the firing rate and the synchronization score (Refer method section 4-3-2). The firing rate refers to the number of spikes per unit time [spikes/sec], and the synchronization score refers to the frequency of events in which other neurons also fire during a certain time window after one neuron fires.

Because the training and test data are originally the same time series, this case is relatively easy to generate and predict. Therefore, it was expected that it would perform close to the best prediction performance when the training and test data are cut out from the same time series.

As a result, Fig. 3-(c) and Fig. 3-(d) show scatter plots between predicted and measured values for both the firing rate and the synchronization score, respectively. In these scatter plots, we observed a concentrated point on the diagonal for both the firing rate and the synchronization score, indicating that the generation was successful [Fig. 3-(c)(d)].

Fig. 3-(g) plots the correlation coefficients of the Firing rate for each brain region used in the measurement. In all regions, the correlation values exceed 0.9, indicating high predictive success, with the exception in RFV [refer to supplemental material].

For further evaluation of the synchronization score, the first and third quadrants of the scatter plot were extracted and histograms were drawn in the direction of the rotation axis. Examples are shown in fig. 3-(e),(f). As seen in these results, it can be observed that the generation is successful as peaks.

From these results, it was found that when the training data and the generated data are obtained from the same region, both the firing rate and synchronization could be nicely reproduced (figs. 3-(g-j)).

In the next section, we will observe the case where the training data and the generated data are obtained from different regions. The results given in this section provided us the approximate values of every prediction performance in the relatively easy problem of generating from training to test data cut out from the same time series (figs. 3-(g-j)). The performance would give us perspective on the highest value when generating across different data shown in the next subsection.

### 2-3. Generation with all brain regions as source and target regions

While training and generating evaluations were performed on data acquired from the same brain region in 2-2, in this section, we also analyze and evaluate the training and target data recorded respectively from two different brain regions groups included in the same one of two datasets [Fig. 4-(a,b)]. The prediction performance was then shown as the average of dataset 1 and dataset 2 [Fig. 4-(g-k)].

**Figure4:**
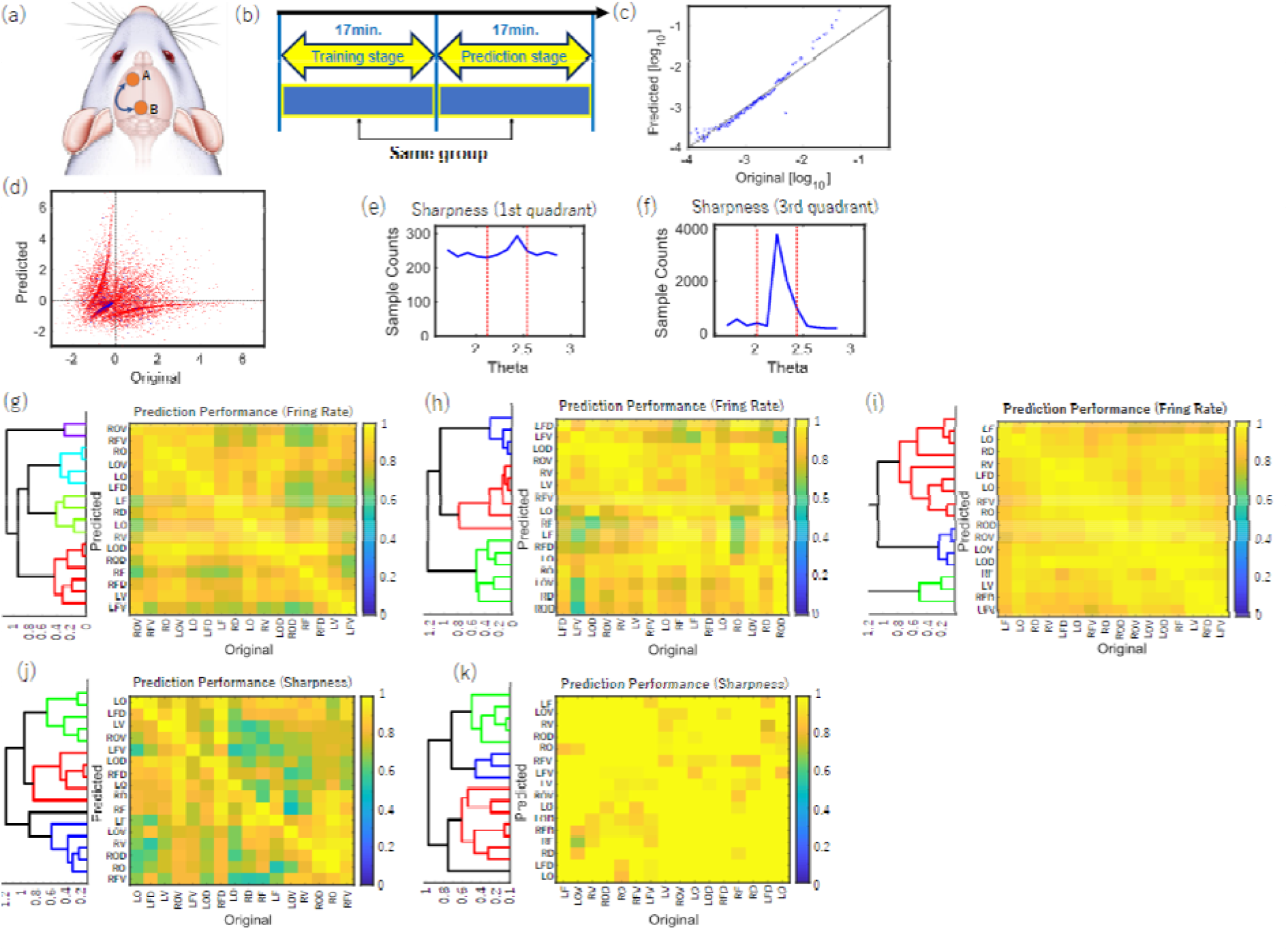
(a) In this evaluation, training data and prediction, generation, data are also prepared from different regions. (b) In the case of (a), training data (first 17 minutes) is obtained from region A and test data (second 17 minutes) from region B. (c) This panel shows the result of predicting the firing rate in a particular case, where the x-axis shows the firing rate in the original test data and the y-axis shows the firing rate in the generated test data by training Multi-layer LSTM. (Again, the x-axis is the synchronization score in the original test data, and the y-axis is the Synchronization score in the generated test data. This two-dimensional distribution was expressed in terms of r-θ rotational coordinates, and panel (e) depicted the density distribution of the number of data in 0-π/2 with respect to θ, i.e., in the first quadrant. Finally, panel (f) is the density distribution of the number of data in π-3/2π with respect to θ, or the third quadrant. In particular, we have confirmed that the distribution is restricted to the third quadrant when the output is from inhibitory cells. The sharpness of the peaks in these histograms (e) and (f) was evaluated by sharpness. (g) is the correlations between generated and truth values in firing rates for all cells, (h) for inhibitory cells, and (i) for excitatory cells. x-axis is the region index of the original data and y-axis is the region index of the predicted data. The correlations of firing rates in all those pairs are plotted as color maps. In addition, these color maps are sorted based on hierarchical clustering. (j) and (k) are the color maps at sharpness in the first and third quadrants, respectively. The point that the sorting is based on hierarchical clustering is the same as in the case of the color map of firing rates.

By comparing between different data, the similarity of their neural activity can be assessed by predictability by generating process rather than by cross-correlation. It should also be noted that in this in vitro experimental environment, the connections between the brain regions are broken, so the similarity between time series data from two brain regions is by no means produced by the interaction of the two brain regions.

Again, the quality of generation was evaluated using the firing rate (Fig. 4-(g-i)) and the synchronization score (figs. 4-(j, k)) (Refer method section 4-3-2).

First, accuracy with respect to firing rate in generation was evaluated simply by crosscorrelation between the firing rate in the original test data and the firing rate in the generated data (Fig. 4-(c)). Color maps of the correlation values between the firing rates of all neurons (Fig. 4-(g)), inhibitory cells only (Fig. 4-(h)), and excitatory cells only (Fig. 4-(i)) are plotted. Hierarchical clustering was performed to sort brain regions that show similar patterns in terms of prediction performance into close indices.

Second, when evaluating the accuracy with respect to the degree of synchrony in the generation, we used the scatter plot (Fig. 4-(d)) between the synchronization score in the original test data and the synchronization score in the generated data.

Then, in the scatter plot, we evaluated the peakness of the angle-dependent distribution in the first (Fig. 4-(j)) and third (Fig. 4-(k)) quadrants of the data distribution as the sharpness (method 4-3-4). In the synchronization results, the data were sorted by hierarchical clustering so that regions with similar characteristics are close to each other.

In all the results so far, the diagonal components are brighter than in other cases because the generation between the same region shows a high prediction performance. However, at the same time, the generation between different regions also sometimes showed high prediction performance at the same level as the generation from the same region.

In a later section (sections 2-5), we will further analyze how such brain-region pairs of nondiagonal cases, showed similar prediction performances with the diagonal cases are related to each other, based on the relative spatial distances and/or structural connectivity between brain regions.

### 2-4. Relationship with connection strength and relative distance

Finally, in order to study how anatomical “closeness” relates with the unevenness of performance in intergeneration between different brain regions in the results obtained in sections 2-2 and 2-3, we evaluated the results in comparisons to the relative distances between brain regions and the strength of structural connections.

As shown in Figure 5-(a), relative distances were calculated based on the relative angles from the regions of interest selected from 16 regional groups. First, the relative distance in the group of interest was set to zero. Then, within the ipsilateral cortex of the group of interest, the relative distance was incremented by +1 with every one angle difference. However, the completely opposite angles were set to +2 since they are adjacent to each other on the same slice plane. (fig 5-(a)).

**Figure5:**
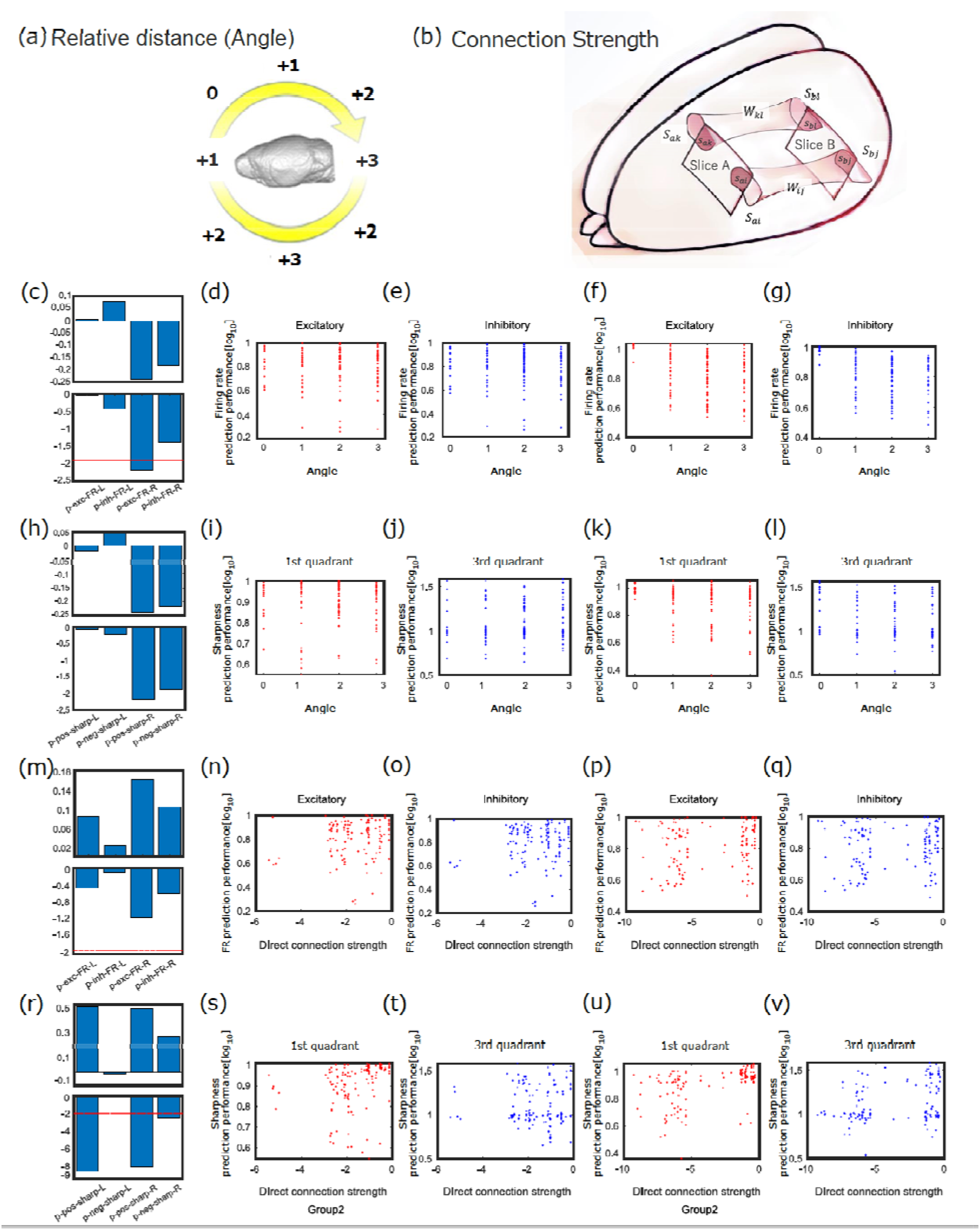
(a) shows the definition of a score calculated based on the relative distance from a certain region of interest (in this case, the Left Frontal Dorsal region). The score at the region of interest was set to zero, and the score was added by one for every shift of one angle from the region of interest. However, the score for the region completely opposite to the region of interest was reduced to +2 because that region is located at the same slice surface as the region of interest and adjacent to the region of interest. (b) illustrates the definition of the values required to calculate the strength of the structural connections between the square recording regions where electrodes were placed. We downloaded the original connection strengths and atlas data from open data shared by Allen institute (https://connectivity.brain-map.org), and reconstructed the connection matrices. In this illustration, the two squares represent the areas (e.g., Slice A, Slice B) where electrical measurements were made. In addition, two examples of connections between them are expressed as pipes. The connection strength between the atlas areas included

We utilized tracer data published by Allen institute with the Mouse Reference atlas for the structural connections [Lein et al. 2007; Dong et al. 2008; Oh et al. 2014]. In this study, we calculated the connection strength between two square recording regions in four steps.: First, we enumerated the cortical areas on the atlas that belonged to the two square recording regions where electrical activity was measured. Second, we calculated the strength of the structural connections between all pairs of the cortical areas belonging to each square region. Third, the connection strength was then normalized by the percentage of area within the region located at either end of the connection. Finally, the normalized connection strength was averaged for all combinations of regions and calculated as the connection strength between electrode recording regions (fig 5-(b)). See Method 4-3-5 for the detailed formula.

Then, we evaluated the relationship between either the relative distance or the connection strength between the electrode recording regions and either the accuracy of predicting the firing rate or the sharpness and accuracy of predicting the synchronization score between the region pairs. This evaluation was performed with the left and right hemispheres separately (fig5-(c-v)). The statistical test here is a Bonferroni correction for a sample size of 4 in the panel.

First, fig.5-(c-l) lists the results for relative distance (angle). For a difference of 1 relative distance (angle) from 0, the predicted firing rate showed a significant difference (fig5-(c-g)) (p=0.006, p<0.01, Bonferroni correction). However, sharpness, which is the prediction performance of connection strength, in relation to relative distance showed no significant trend.

Second, fig5-(m-v) lists the results for the connection strength. No significant trend was observed in the prediction of firing rate for connection strength in any condition (fig5-(m-q)). However, there was a significant positive correlation between connection strength and predicted sharpness in the first hemisphere (Left hemi.: p=5.510^-9^, Right hemi.: p=2.010^-8^, p<0.01, Bonferroni correction), which was common in the left and right hemisphere.

In general, the results indicate that the prediction performance of the index of firing activity in the multi-layer LSTM is related to both the relative distances between measurement sites and the strength of the structural connections. We will address more in-depth discussions of this relationship in the discussion section.

in the square region of the electrical recording was calculated for each connection as a normalized value in terms of the percentage of the area, expressed as a dark red area in fig. 5-b, of the intersection of the region of the electrical recording with the area of the atlas at both ends of the connection (see Method III-6). The normalized value as a percentage of the area, expressed as a dark red area in fig. 5-b, was calculated for individual connections (Refer to Method III-6 for details).

In the square region of the electrical recording, the normalized value, expressed as a dark red area, was calculated for each connection as a percentage of the area of the intersection of the region of the electrical recording with the area of the atlas at both ends of the connection (see Method III-6). The normalized value as a percentage of the area was calculated for individual connections (Refer to Method III-6 for details).

Then, the connection strength between the regions of electrical recording was calculated as an averaged quantity of connection strength in all connected pairs of atlas areas at both ends.

Panels (c) ~ (l) plot the prediction results for the relative distances (angles) between the measurement regions. Among them, panels (d) ~ (g) plot scatter plots with the predicted firing rates for the relative angles between the regions, and panel (c) summarizes the correlation values (upper panel) and p-values (lower panel) for the points between Angle=0 and Angle=1 in panels (d) ~ (g) as a four-bar graph corresponding to the order from (d) to (g). Among (d)-(g), (d) represents results to excitatory neurons in the left hemisphere, (e) to inhibitory neurons in the left hemisphere, (f) to excitatory neurons in the right hemisphere, and (g) to inhibitory neurons in the right hemisphere.

Panels (i) through (l) are plotted as scatter plots with sharpness, prediction performance of Synchronization score, as the vertical axis relative to the relative distances (angles) between the measurement regions. The difference in the meaning of the x-axis in the four panels (i) ~ (l) means results to the first quadrant of the left hemisphere, (j) to the third quadrant of the left hemisphere, (k) to the first quadrant of the right hemisphere, and (l) to the third quadrant of the right hemisphere. In the panel (h), the correlation values (upper panel) and p-values (lower panel) for the points at Angle=0 and Angle=1 in (i)~(l) are summarized as bar graphs. The panels (n)~(q) and (s)~(v) are the same as the panels (d)~(g) and (i)~(l), except that the horizontal axis is the connection strength. Then, the correlation values in the upper panel and p-values in the (lower panel at (n)~(q) are summarized in (m), and the correlation values (upper panel) and p-values (lower panel) at (s)~(v) are summarized in panel (r). The significance level is set at about p=0.01 and the dotted lines are overlaid, and it can be read that the p-values corresponding to (s) and (v) are much lower than that level.

## 3. Discussion

### 3-1. Summary of the results

In this study, we developed a new approach to evaluate the homology between two regions through the generation and evaluation of synthetic neural spike data using a Multi-layer LSTM network.

Specifically, spiking data for a group of over 100 neurons were measured from slices taken from 16 cortical regions of the mouse, and from those 16 regions, 16×16=196 different pairs of training and test data were prepared.

When interpreting the given results, it is important to keep in mind that in slices cut from one region, connections to other regions are physically severed. It was well expected that our newly applied spike generation technique between different regions would not work at all because the recording regions are disconnected from each other.

However, surprisingly, the results showed that there are hidden rules in the spike data that allow the Deep Neural Networks used in this study to even generate complex spike sequences to the point of reproducing them with nontrivial accuracy. It is extremely difficult for the human eye to decipher the rules utilized in their generation.

It should also be emphasized that even if one creates a detailed computational model of neural systems, it is actually very difficult to prepare a computational model that generates spikes that somehow reproduce the synchrony among the many pairs of neurons in the system [Nolte et al., 2019; Dura-Bernal et al., 2019; Feldotto et al., 2022]. The findings of this study can be summarized in the following three categories:

First, the case of learning and generation among time series of different time periods in the same data showed clearly significant prediction performance, not only in terms of firing rate, but also in terms of the degree of synchrony.

Second, even in the predicted performance among the regions measured from different regions, surprisingly, there were some combinations that came close to the performance for the same data.

This indicates that the characteristics of electrical activity within cortical local circuits have enough commonality or universality to generate each other even if the regions are different. There is no precedent for showing this commonality through the mutual generation of activity.

Third, we compared the prediction performance of firing rate and synchronization with the relative distance (angle) between the measured regions and the strength of the structural connections. The results showed that there was a significant difference in the prediction performance of firing rates between generations made between the same region and those made between regions that were one relative distance (angle) adjacent to each other. Moreover, and more surprisingly, although no significant correlation was observed between structural connection strength and prediction performance of firing rate, significant correlations between structural connection strengths and sharpness, which is prediction performance of synchronization score, were observed in both left and right hemispheres or in one hemisphere in some cases.

### 3-2. Potential contributions of data generation technology to improve the 3Rs

The methods developed in this study have very important significance for the fundamentals of animal experimentation. In neurophysiological experiments, synchronization still plays an important role in quantifying neural representations of neural interactions and cognitive functions [Hunt et al., 2018; Galske et al., 2019; Knoblich et al., 2019; Luo et al., 2022]. If such spiking data can be generated from artificial models with high accuracy, there is no need for redundant experiments. As a result, they can contribute to the 3R principle, Replacement, Reduction and Refinement, regarding animal experiments [Törnqvist et al. 2014]. Such techniques will become even more important for rare data, where large amounts of data are difficult to obtain [Alexandra, 2020; Perretta, 2009]. Therefore, it is likely that research schemes to quantify the similarity of different datasets measured under different conditions in terms of their inter-generational capabilities will expand in the future.

### 3-3. Interpretation of non-uniformity in generation performance

It is also important to keep a dispassionate attitude in considering high projection performance.

For example, in the relation between the prediction performance of the firing rate and the relative distance (angle), there was a significant difference in the prediction performance of the firing rate between the case of generations made between the same regions and the case of generations made between one adjacent region. Remember, however, that there is a possibility that the advantage of being cut from the same data (beyond being in the same region) worked in the prediction between the same regions. Therefore, further verification is necessary to check if it is not an artifact. Nevertheless, the trend in the first quadrant that synchronization score increased with each increase in structural connectivity is a non-trivial trend that cannot be explained by such reasoning.

### 3-4. Argumentation on structural connectivity

It should be noted that there have been numerous previous studies on structural connectivity. First, the hierarchy of information processing, which is described in terms of structural connectivity patterns between cortical areas, has been logically defined based on differences in laminar patterns along the cortical depth direction. This is the definition conventionally employed in the analysis of wide-ranging structural connectivity patterns, mainly for the visual system [Fellman, VanEssen, 1991].

Some researchers have attempted to understand the hierarchy of information processing by integrating this pattern with the auditory and somatomotor systems in informatic ways [Modha et al., 2010]. The understanding of the hierarchical structure of information processing was then extended to an understanding of hierarchy in the sense of going from peripheral areas closely connected to the peripheral nervous system of information processing to central core areas corresponding to the association cortex [Kötter, Wanke, 2005].

We can also confirm that the hierarchy of information processing reflected in connection structure patterns is related to cell density [Shimono, 2013]. Furthermore, as we obtained better systematic data on connection strength, it became clear that there is a clear empirical relationship between connection strength and spatial distance [Markov et al., 2014].

Various studies have reported that the pattern of structural connections is similar to the pattern of functional connections defined on the basis of synchronization of activity between joined brain regions [Honey et al. 2007, 2009; Mišić et al. 2016].

### 3-5. Prospects and future challenges

#### 3-5-1. Comparisons of generation performance with physiological indicators

Even with the important background knowledge described in the above subsections, it is quite surprising that we observed a significant positive correlation between structural connection strength and sharpness for the first quadrant (for example, fig. 5-s), since we measured each brain region as slices. This is because, as we have mentioned many times, we measured neural activity from brain regions after sectioning them as slices.

Structural connectivity patterns in mice have been measured and analyzed on a large scale with increased resolution in a way that also integrates with genemics or transcriptomics studies [Lein et al., 2007; Harris, et al., 2019; Hawrylycz et al., 2015]. This study also aided the structural wiring pattern information obtained in those studies [Oh, et al., 2014].

Since such genes and transcription factors are internalized characteristics of each brain region, there is a possibility that they are related to the ease of mutual generation of neural activity in each brain region, which we have discovered. Therefore, it is a future challenge to investigate the relationship between these genomics and transcriptomics and the internal characteristics of activities within individual brain regions.

#### 3-5-2. Improvements of generation and evaluation methods

One of the other issues is the improvement of the generation method using multilayer LSTMs and the evaluation method. When generation does not work well, the disappearance of the diagonal component in the scatterplot of the relationship between the predicted and correct answers generally occurs. Although this report does not go into detail, several characteristic patterns were observed following the disappearance of the diagonal component in the scatter plots. By classifying these characteristic patterns and exploring their causes individually, guidelines for their generation and evaluation at high performance will be more mature than now.

In this study, we chose the region that brings together the cell groups as the square recording region in the electrical measurements. In other words, the brain regions of the atlas provided by the Allen institute were grouped together within the square measurement area, and the analysis was performed to compare them with the structural connections. Another future task is to analyze cell groups separately according to the atlas brain area segmentation provided by the Allen institute. This will allow for a more stable combination of cell groups, including those within a single region, which will positively affect the generation and prediction results. -This could have a positive effect on prediction performance. However, new ideas are needed to deal with the fact that the number of cells in each region may differ due to the difference in size of each region.

This study collectively analyzed populations of neurons existing in the square region used for electrical measurements. However, there may be cases where histologically distinct brain areas coexist within these square recording regions. Therefore, one of the future tasks is to analyze the neuron groups according to segmentations of brain atlases such as the Allen brain atlas. This will allow us to include only neuron groups within a single region, and to make the groups of neurons belonging to each individual group more uniform. As a result, we expect that this will have a positive influence on the performance of the generation and prediction. However, new ideas are necessary because the number of cells within a region should be different due to differences in sizes of these regions.

Although new models do not necessarily improve performance, changing the learning-generating model from a multi-layer LSTM to a model such as Transformer [Vaswani et al., 2017] also has the potential to contribute to improved performance.

### 4. Final remarks

This study showed that the activity of multiple neuron groups in the cortex can sometimes be generated reciprocally, even between different regions. We also showed that the non-uniformity of the reciprocal generation can be explained, to some extent, by the relative positional relationships and structural wiring. In such a method, the combination of the original data and the target data to be generated is very broad.

In the future, as the physiological interpretation is deepened and the prediction performance is improved, it will become the basis of a method to “measure” physiological data through mathematical models instead of experiments, and is expected to contribute to the comparison of experimental data among animal species and the performance evaluation of model animals as well as the 3Rs. This method is expected to contribute not only to the 3Rs but also to the comparison of experimental data among animal species and the evaluation of model animal performance.

In this study, we demonstrated for the first time that the activity of groups of multiple neurons in the cerebral cortex can often be mutually generated, even between different brain regions. We also suggested that the non-uniformity of performance in mutual generation can be somehow explained by relative spatial positions and structural connections.

It should be emphasized that such a generation method allows a very wide range of choices in what data to use for the combination of original data and target data to be generated. In other words, this method allows us to evaluate similarities between a wide range of neural activities.

## 4. Methods

### 4-1. Data acquisition of neuronal activities

We used neuronal spike data recorded and studied in detail in our past study. Here, we briefly explain the experimental procedure utilized in the past study [Kajiwara et al., 2021; Shirakami, 2021, Matsuda et al., 2022]. The whole experimental processes are also now open in a video journal [Ide et al., 2019].

We used female C57BL/6J mice (n=32=16×2, aged 3-5 weeks). This study grouped the cortex (mainly the neocortex) into 16 groups, and prepared two sets of data for each of these groups (Fig. 1-(b), supplemental material). All animal procedures were conducted in accordance with the guidelines of animal experiments of Kyoto University (KU), and have been approved by the KU Animal Committee.

This study recorded neuronal spikes from cortical slices with a Multi-electrode array(MEA) system (Maxwell Biosystem, MaxOne) with refluxing an artificial cerebrospinal fluid (ACSF) solution that was saturated with 95% O_2_/5% CO_2_ [Kajiwara et al., 2021; Ide et al., 2019].

Prior to slice preparation, mice are thoroughly anesthetized (1%-1.5% isoflurane), cervical vertebrae was dislocated, and brains are removed. We immersed the removed brain in a cutting solution, an ice-cold solution used to prevent brain deterioration, and bubbled it with oxygen continuously. The brain was cut with a vibratome (NLS-MT, DOSAKA EM CO.,LTD), scanned, and sliced by a slicer in the target region of the brain. 300-μm slices were made by changing the height of the cutter. The slices were immersed in an ACSF solution, saturated with 95% O_2_/5% CO_2_, for one hour before electrical measurements were taken.

The recording area of the MEA used was 3 x 4mm^2^, and 26,000 electrodes were uniformly arranged at 15-μm intervals (on the order of cell spacing distance). This high-density electrode arrangement enables accurate determination of neuron location. For the main measurement, we used on the 1020 electrodes, selected, as receiving strong input from the neurons, in the 20-minute prescan.

We performed spike sorting (Spyking Circus software) from the time series obtained in the main measurement and converted to time series binary data of the activity of the cell population. Refer these following references about the details of the experimental procedure [Ide et al., 2019, Kajiwara et al., 2021; Shirakami et al., 2021, Matsuda et al., 2022].

### 4-2. Arrangements of data formats to input into the analytical model

The spike data can be represented as a binary matrix X_ti_ where t and i are time and neuron indices: if neuron i is firing at time t, then X_ti_= 1, and otherwise X_ti_= 0. We sorted the neurons so that inhibitory neurons have earlier indices than excitatory neurons. Within each set of neurons, neurons are sorted in the order of layers (from 6 to 1). Each segment of spike data was split into training and test data. After removing the first 30 minutes, two 17-minute-long segments, earlier segment (30–47min) and later segment (48–65 min), were cut out.In the same region cases, we used the earlier segment as training data and the later segment as test data. In contrast, in the different region cases, one of the earlier segments from the two regions was used as training and the other one as test data.

### 4-3. Analysis

#### 4-3-1. How to connect different data ?

In this study, we use a Recurrent Neural Network (RNN) with long short-term memory (LSTM) units for generating spike data. [Hochreiter, Dchmidhuber, 1997]. An RNN takes time series data as input and outputs time series data of the same length, and it iteratively processes a time-sliced (vector) data at a time instant. Thanks to its design, the information accumulated from the past data is to be utilized along with the current input.

Simple RNNs have difficulty for dealing with long-range dependence and have limited ability to learn the influence of data from the distant past. The LSTM units are proposed to alleviate this problem by introducing an architecture to deal with long-term memory. An LSTM unit contains three gates: “input”, “output” and “forgetting”. These gates determine the degree to which information is allowed to pass through depending on the conditions. The LSTM unit can store information more efficiently by gradually changing the long-term memory while maintaining the RNN structure itself.

Here, let us describe the history of LSTM. In 1997, Hochreiter and Schmidhuber et al. proposed LSTMs with cells and input and output gates, and in 1999 Gers et al. introduced an oblivious gate in the LSTM structure. In 1999, Gers et al. introduced an oblivion gate in the LSTM structure, which allows the LSTM itself to reset its own state. In 2000, Cummins et al. added peephole connections to allow cell-to-gate coupling; in 2014, Cho et al. proposed a gated regression unit, and a subsequent speech recognition using LSTM showed a 95.1% recognition accuracy. The LSTM network has been applied to speech recognition [Graves, et al. 2013], handwriting generation [Graves, 2013], language modeling [Sundermeyer et al. 2012], and many other tasks. The model is still widely used today.

We utilize a multi-layer LSTM network, which can learn even longer-range dependence on the data than a single-layer LSTM network can (Fig. 6). The following hyperparameters of the LSTM network, which are different from the parameters to be trained from data, were used in this study.

**Figure 6.**
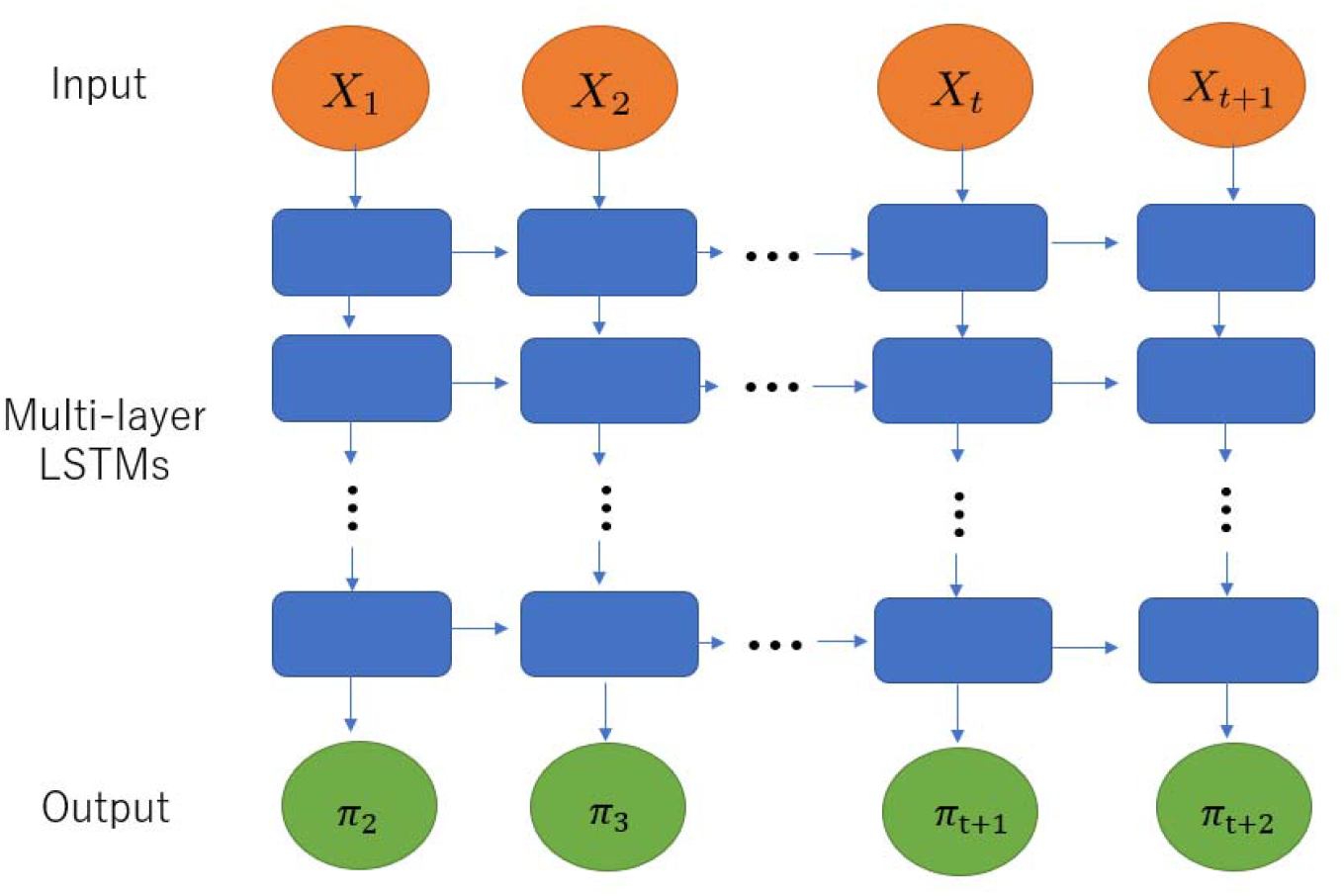
This study utilizes multi-layer LSTMs (multi-layer LSTMs), which are networks of LSTMs layered on top of each other to allow for even longer-term learning than single-layer LSTMs. The input is a prediction of the probability of firing, and the output is the result of the target prediction. Whether or not a target has fired is determined by whether or not the measurement results exceed a threshold value.

First, the number of layers in the LSTM network is five, including the input and output layers. The number of LSTM units, which is the dimension of data in the intermediate layers, was 128 (same for all intermediate layers). The dimension of the input and output vectors, which are the number of neurons, was also 128.

The input of our LSTM network at time is the vector =( ), and the output at time is the vector of probabilities that neurons will be firing at (each element takes a real value between 0 and 1).

The network is trained with a loss function defined in the next section.

After training, to generate spike data, we use the test data as input and apply thresholding to the output : neuron is regarded to be firing at time if >, where is a threshold value.

#### 4-3-2. Loss function

The binary cross entropy loss function is often used for training a neural network for data generation.

It is formulated as where πti denotes the network’s output and Xti denotes the training data.

In the spike data, most elements in X_ti_ are 0 since the firing rates are small. In such a situation, the binary cross entropy loss often makes the network learn to output very small firing probabilities, which hinder the appropriate generation of spike data. To deal with such imbalances between states, a refined loss function called focal loss has been proposed [Lin et al., 2017].

The loss function has an additional parameter γ and is given as

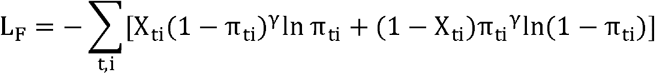

Here, the parameter γ>0 induces the two terms to balance since the non-firing probabilities 1-π_ti_ are much larger than the firing probabilities π_ti_.

We use the value γ=2 as suggested in the original study [Lin et al., 2017].

The number of epochs for training was set to 350. We chose this number because, although the loss value had converged in about 150 epochs, the prediction performance of the firing rate and the reproduction performance of synchronous firing improved as the training further proceeded. The Adam optimizer [Kingma, Ba, 2014] was used for training. The batch size, which is the number of data segments used in one update of the training process, was set to 64.

#### 4-3-3. Evaluation of similarity between generated and real data

We analyzed the probability of synchronous firing between neurons to evaluate generated data with respect to the reproducibility of the property that the timings of neuron firing are synchronized between two cells with a specific delay. To formulate an evaluation metric, suppose a situation where after neuron i fires neuron j fires with an acceptable delay D. If the firing of neuron i has a positive effect on the firing of neuron j, then we expect a larger firing probability of neuron j within some acceptable delay D than its mean firing probability (i.e. firing rate).

Let us express the conditional probability that neuron j fires at least once within delay D after neuron i fires as q(j|i; D). If neuron i and neuron j are not synchronous (i.e. they are independent), the conditional probability is given as

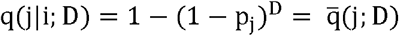

where p_j_ expresses the firing rate of neuron j. We used the following indicator Z to evaluate how much the actual conditional probability q(j|i; D) is biased from the null hypothesis 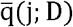:

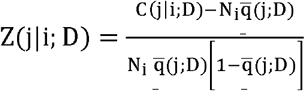

where N_i_ is the number of firings of neuron i, C(j|i;D)=Nj(j|i;D) is the number of times that neuron j fires at least once within delay D after neuron i fires, and the denominator represents the standard deviation in the null hypothesis.

We call this quantity synchronization score. We obtained a scatter plot of the synchronization score plotted with the real data on the horizontal axis and the generated data on the vertical axis (refer fig. 3-(d),4-(d)), and a histogram with the rotation angle θ from 0 degrees as the main axis.

The histogram is called sharpness positive (refer fig. 3-(e),4-(e)), which is calculated as the ratio of the area around the peak (width of π/4) in the first quadrant (Θ = 0 ~π/2) to the area at other angles included in θ = 0~π/2. The sharpness of the area around the peak (width of π/4) in the third quadrant (θ = π~3π/2) is also calculated in the same way and is called sharpness negative (refer fig. 3-(f),4-(f)).

Just before detecting those peaks, we performed a linear regression on the histogram, and only the trend of the slope of the line was removed. The acceptable delay D was set to 1. The reason is that the sharpness for other values of D had a negative effect on observing the relationship between the generated and real data, blurring the diagonal components of the scatterplot than when D=1.

#### 4-3-5. Definition of angles between recording regions

The angles between regions were classified into eight groups in 45-degree increments in the direction of rotation with the line connecting the left and right ears as the axis. In other words, the left and right hemispheres were taken into account and classified into 8 × 2 = 16 groups (fig.5-(a)). Note that in this study, the angle between two groups is also called the distance between two groups. 16 groups are named as shown in Table 1, with two data belonging to each region (Refer the supplemental material).

#### 4-3-6. Definition of connection strength

In this study, we superimposed the cut-out position of each slice on the Allen reference atlas (ver.3, https://mouse.brain-map.org/static/atlas) and extracted the brain region in the Allen atlas that each slice includes.

Now, as shown in Fig. 5-(b), we consider two slices and call them slice A and slice B.

Then, focusing on brain regions a_1_, a_2_, …, a_n_ included in slice A (divided by the atlas) and brain regions b_1_, b_2_, …, b_m_ included in slice B, We obtained the connection strength W_ij_ between ai (i = l~n) and b_j_ (j = l~m) for all pairs.

Next, calculated the ratio R_ai_ = s_ai_/S_ai_ (i = l~n) of the area S_ai_ of the entire brain region on the slice A cross-section to the area s_ai_ that is in the recording region on the slice A. Similarly, R_bj_ (j = l~m) is calculated for slice B. Then, for example, W_ij_R_ai_R_bj_was calculated for a connection pair of regions i and j.

Finally, we obtained the connection strength between slice A and slice B regions by adding it between all i and j, 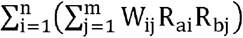.

## Acknowledgement

MS has been supported by several MEXT Grant-in-Aid for Scientific Research (B) (20H04257, 21H01352) and the Leading Initiative for Excellent Young Researchers (LEADER) program. EN is supported by several MEXT Grant-in-Aid for Scientific Research (B) (22H03661) and Scientific Research (C) (21K02846, 21K12187). The imaging of MRI and immunostaining of this work were performed in the Division for Small Animal MRI, Medical Research Support Center, Graduate School of Medicine, Kyoto University, Japan. We acknowledge Doris Zakian excellent comments to edit this manuscript, and all support from Innovative Support Alliance for Life Science, and the Hakubi Center in Kyoto university to complete this study.

